# The transcriptional landscape of a hepatoma cell line grown on scaffolds of extracellular matrix proteins

**DOI:** 10.1101/2020.07.18.191395

**Authors:** Souvik Ghosh, Anastasiya Börsch, Mihaela Zavolan

## Abstract

The behavior of cells *in vivo* is complex and highly dynamic, as it results from an interplay between components of the intercellular matrix proteins with surface receptor and other microenvironmental cues. Although the effects of the cellular niche have been investigated for a number of cell types using different molecular approaches, comprehensive assessments of how the global transcriptome responds to 3D scaffolds composed of various extracellular matrix (ECM) constituents are still lacking. In this study, we explored the effect of the biomechanical parameters of Collagen I and Matrigel (ECM) on transcriptional gene regulation in a cell culture system. Using Huh-7 cells cultured on traditional cell culture plates or on the components of the ECM at different concentrations to modulate microenvironment properties, we have generated transcriptome sequencing data that may be further explored to understand the differentiation and growth potential of this cell for the development of 3D cultures. Assessment of the hepatocyte phenotype in relation to our transcriptomic data set would be very useful for the development of systems mimicking the *in vivo* structure and function of liver cells which still remains a challenge.

## Introduction

The liver is a critical hub for numerous physiological processes. These include macronutrient metabolism, blood volume regulation, immune system support, endocrine control of growth signaling pathways, lipid and cholesterol homeostasis, and the breakdown of xenobiotic compounds, including many current drugs^1^. It is composed of about 80% hepatocytes and 20% non-parenchymal cells such as stellate cells, sinusoidal endothelial cells, and Kupffer cells^1^. Isolated hepatocytes generally cultured in two-dimensional (2D) culture using polystyrene-coated culture flasks lose their native morphology, polarity and functionality, which subsequently limits their effectiveness in applications such as toxicity screening of drug metabolites^2^. It has been reported though, that culturing hepatocytes between collagen layers, could be a better alternative for primary hepatocyte culture ^3,4^. Such three-dimensional (3D) cultures for primary hepatocytes proved to be better in maintaining hepatocyte phenotype and cell polarization.

The Extracellular matrix (ECM) is recognized as a dynamic structure that provides a supportive scaffold and actively regulates biological functions of cells, at least partly by the interaction with specific cell surface molecules ^5^. Three-dimensional matrices that are used to study cell behavior in a tissue-like environment in normal and pathological conditions are often made primarily of collagen, amongst all other components of the ECM ^6,7^.

Development of a culture system mimicking the *in vivo* structure and functions of liver cells still remains a challenge. We sought to contribute to resolving this challenge by setting up a cell line-based experimental system that exhibits hepatocyte-like polarity. The system is based on Huh-7, a well-established, differentiated hepatocyte cell line that was generated in 1982, from a liver carcinoma of a 57-year old Japanese male^8^. The ability to propagate Huh-7 cells in media containing defined components broadens the relevance of this cell line from a model of oncogenesis to one also suitable for the elucidation of regulatory mechanisms of gene expression. The properties of these hepatoma cells allow systematic studies of *in vitro* effects of various compounds on their growth and metabolism^9^. An additional benefit of using the Huh-7 cell line was its permissiveness to Hepatitis C virus genomic replication.

Pathophysiological changes / regeneration of the liver is accompanied by the remodelling of the ECM ^10–12^. To gain molecular insight into the changes that contribute to the phenotype, we developed 3D growth models of Huh-7 on different concentrations of Collagen I matrix. Furthermore, in line with previous studies, we also used Matrigel as a basement membrane matrix in several concentrations. Matrigel is a gelatinous protein mixture secreted by the Engelbreth–Holm–Swarm mouse sarcoma cells that resembles the complex extracellular environment found in many tissues and is used to produce thick 3D gels for cell culture ^13^. It has previously also been documented that Matrigel-cultured Huh-7 cells assembled into 3D spheroids, whereas standard 2D-cultured cells formed the typical epithelial monolayer ^14^. The cues provided by the niche, are known to be important for modulating cellular behaviors such as differentiation, viability, and proliferation. Being able to modulate these behaviors via simple changes in the concentration of matrix components would be very convenient for smart scaffold designs. In the current study, we identify several transcription factors that modulate the gene expression profile of the cells. Furthermore, we also observe that several of these factors change linearly with changes in concentration of matrices used for growth which implies robust roles of these factors and their involved pathways. Interestingly, analysis of the data sets also reveal the great heterogeneity in the incurred changes between the two different types of matrices used to seed growth. Our data also reveals overall changes in critical hepatic functions, like insulin secretion, drug metabolism, amino acid catabolism highlighting the roles of extracellular matrix components on liver function. This comprehensive analysis of the gene expression profiles and the corresponding transcription factor activities would be extremely useful for futuristic design of matrix scaffolds for three dimensional growth of cells/ organoids derived from liver to mimic real life conditions.

## Methods

### Coating of plates with collagen and matrigel matrix

#### Collagen Coating

Collagen coating of plates was performed inside a cell culture hood to prepare tissue culture plates for seeding cells on a collagen matrix. Initially, Collagen (Thermo Fisher Scientific, Catalogue #A1048301), sterile 10X phosphate buffered saline (Thermo Fisher Scientific, Catalogue #AM9625), sterile distilled water (dH2O), and sterile 1N NaOH (Merck, Catalogue # S2770-100ML) was thawed on ice. For our experiment we used final Collagen concentration of 1mg/ml and 0.5 mg/ml. To prepare a total of 2 ml of solution of collagen of each concentration we used the following formula.

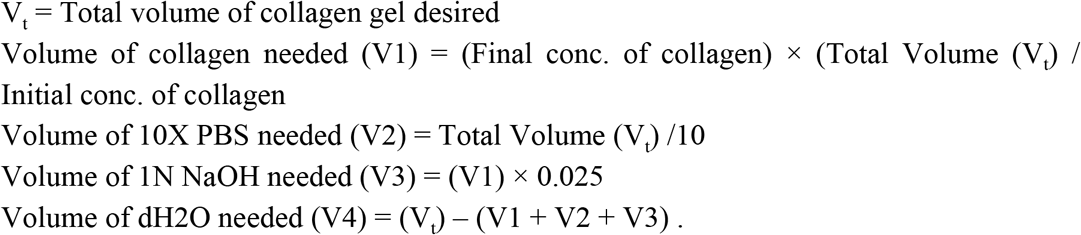

Subsequently, in a sterile tube, dH2O, 1N NaOH, and 10X PBS were mixed. The estimated amount of collagen was slowly pipeted into the tube, with mutiple rounds of pipetting the solution up and down to achieve a homogeneously mixed solution. The resulting mixture was checked to ensure that it achieved a pH of 6.5–7.5 (optimal pH is 7.0). The diluted collagen solution was then dispensed carefully into the desired plates (500 μl per well of a standard 6 well tissue culture plates), on ice. The ice was used to ensure uniform coating of the plate with the cold temperature thwarting rapid gelling at room temperature. The plates were then incubated at 37°C in a humidified incubator for 30–40 minutes until a firm gel was visually observed. Lastly, the wells were rinsed twice with 1X PBS and cell culture medium before seeding of cells.

#### Matrigel® Growth Factor Reduced (GFR) Basement Membrane Matrix Coating

To prepare tissue culture plates for Matrigel (Corning, Catalogue # 354230) coating, which would be used for seeding cells, the matrigel matrix was thawed before use overnight by submerging the vial in a 4°C refrigerator. Once the Matrigel matrix was thawed, the vial was swirled several times on ice to ensure even dispersion of material. We opted for three different concentrations of matrigel (1.5 mg/ml, 3.0 mg/ml and 6.0 mg/ml) for coating the cell culture plates. The stock matrigel solution was diluted with ice cold Opti-MEM I Reduced Serum Media (Thermo Fisher Scientific, Catalogue # 11058021). The solutions were thoroughly mixed and then kept on ice until further use. Additionally, cell culture plates, tip boxes to be used were also pre-conditioned in a 4°C refrigerator before use. The diluted Matrigel solutions (500 μl) prepared earlier were dispensed into each well of a 6 well plate in triplicates for each dilution. The culture plates were transferred to a 37°C humidified incubator for gelling of the matrigel solutions for at least 30 minutes or until visible gelling was observed. The wells were rinsed with 1X PBS and cell culture medium before seeding of cells.

#### Cell Culture and Imaging

Huh-7 cells grown in cell culture flasks were washed with PBS, and then trypsinized with Trypsin-EDTA (0.05%), phenol red (Thermo Fisher Scientific, Catalogue # 25300054). Following completion of cell detachment from the flask, growth media (DMEM high glucose (Sigma, Catalogue # D6546) supplemented with 2mM L-Glutamine (ThermoFisher Scientific, Catalogue #25030081) and 10% FCS (Amimed, Catalogue #2-01F00-I) was added to stop the process. Subsequently the cells were spun in a centifure at 200g in a swinging bucket rotor and washed twice with PBS. Finally the pellet of cells were suspended in growth media at a concentration of 5000 cells/μl. 25000 cells were then used to seed each well of a 6 well plate with or without the basement matrices as prepared earlier. Cells were observed every day, with growth media being changed every alternate day for a total duration of a week. At 1 day and 7 days post seeding, cells were imaged with an Olympus Phase Contrast microscope, and live snapshots were taken with an Olympus CKX41 inverted microscope equipped with a SC 30 digital camera (Olympus). The images were procured on an inverted phase contrast microscope with Cell Sens software. All images documented were then processed with OMERO (an initiative of the open microscopy environment https://www.openmicroscopy.org/) for digital zoom.

### RNA Seq

#### Sample preparation

Total RNA was quality-checked on the Bioanalyzer instrument (Agilent Technologies, Santa Clara, CA, USA) using the RNA 6000 Nano Chip (Agilent, Cat# 5067-1511) and quantified by Spectrophotometry using the NanoDrop ND-1000 Instrument (NanoDrop Technologies, Wilmington, DE, USA) and adjusted to a concentration of 20ng/μl. 1µg total RNA was used for library preparation with the TruSeq Stranded mRNA Library Prep Kit High Throughput (Cat# RS-122-2103, Illumina, San Diego, CA, USA). Libraries were quality-checked on the Fragment Analyzer (Advanced Analytical, Ames, IA, USA) using the Standard Sensitivity NGS Fragment Analysis Kit (Cat# DNF-473, Advanced Analytical). Samples were pooled to equal molarity. Each pool was quantified by PicoGreen Fluorometric measurement to be adjusted to 1.8pM and used for clustering on the NextSeq 500 instrument (Illumina). Samples were sequenced using the NextSeq 500 High Output Kit 75-cycles (Illumina, Cat# FC-404-1005). Primary data analysis was performed with the Illumina RTA version 2.4.11 and base calling software version bcl2fastq-2.20.0.422.

#### RNA-Seq data processing

RNA-Seq reads were subjected to 3’ adapter (5’-AGATCGGAAGAGCACACGTC-3’) and poly(T) trimming at the 5’ end using Cutadapt v1.9.1 ^15^. Trimming of poly(T) stretches at the 5’ end was performed, because the RNA-seq reads originated from the antisense strand. Reads shorter than 30 nucleotides were discarded. As the reference human transcriptome, we considered sequences of protein-coding transcripts with support level 1-3 based on genome assembly GRCh38 (release 96) and transcript annotation from Ensembl database ^16^. The kallisto v0.43.1 software was used to assign the filtered reads to transcripts ^17^. The default options of kallisto were utilized for building the transcriptome index. For aligning single-end RNA-Seq reads we used the options “--single” and “-l” and “-s”, the latter corresponding to the mean and standard deviation of fragment length, respectively, estimated for each sample with the BioAnalyzer instrument. Since reads originated from the antisense strand, the option “--rf-stranded” was used. The option “--pseudobam” was used to save kallisto pseudoalignments to a BAM file.

Mapped reads were then assigned to transcripts in a weighted manner: if a read was uniquely mapped to a transcript, then the transcript’s read count was incremented by 1; if a read was mapped to *n* different transcripts, each transcript’s read count was incremented by 1/*n*. Trimming the 3’ adapter and poly(T) stretches, indexing the reference transcriptome, mapping the RNA-Seq reads to transcripts and counts of reads assigned to individual transcripts were performed with a Snakemake framework ^18^.

The expression of each transcript *t*_*i*_ was then estimated in units of transcripts per million (TPM) by dividing the read count *c*_*i*_ corresponding to the transcript by the transcript length *l*_*i*_ and normalizing to the library size:

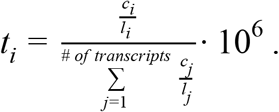

The expression level of a gene was calculated as the sum of normalized expression levels of transcripts associated with the gene.

#### Aligning gene expression with principal components

The gene expression matrix with samples as columns and log2-transformed normalized gene expression in TPM units as rows was centered to make the data comparable both across samples and genes. The centered gene expression matrix was further subjected to the principal component analysis (PCA). For further analyses, we focused on the first two principal components, PC1 and PC2, which explained ~85% of the variance (see Figure 2A,B).

Then we quantified how much individual genes contributed to PC1 and PC2, respectively, and identified genes that induced the difference between samples indicated by these PCs ^19,20^. Figure S1 depicts a toy data set visualizing the concept of the analysis. The toy data set consists of three ‘samples’ and 26 ‘genes’ measured for each sample. Gene expression in these samples was randomly sampled from the multivariate normal distribution. Here, every gene is represented as a vector in the sample space, where rows of the centered gene expression matrix correspond to the coordinates of these vectors. Black vectors indicate PC1 and PC2, respectively, and a blue vector corresponds to a gene vector. For determining the contribution of individual genes to a PC, we calculated the magnitude of the projection of gene vectors on the principal component (PC) (dashed blue line) and the correlation with the PC (cos α). Genes for which the absolute correlation was ≥ 0.6 and the absolute z-score of the projection was ≥ 1.96 were extracted as contributing most to the variance explained by the PC.

#### Gene ontology analysis

To characterize functions of genes contributing most to PC1 and PC2, we performed gene ontology (GO) analysis using Database for Annotation, Visualization and Integrated Discovery (DAVID) ^21^. “GOTERM_BP_DIRECT”, “GOTERM_MF_DIRECT” and “GOTERM_CC_DIRECT” categories were used for the gene annotation. As the reference gene list, we considered genes having the expression of at least 1 TPM in the number of samples corresponding to the minimum number of samples in each condition. We declared a GO term to be significantly enriched if the corresponding p-value was smaller than 0.01. For inspecting protein-protein interactions of genes from top enriched GO terms we used STRING (https://string-db.org/)^22^.

#### Differential expression analysis

Differential expression analysis was performed with EdgeR, available through the Bioconductor package ^23^. Note that a gene was included in the analysis only if it had at least 1 count per million (CPM) in the number of samples corresponding to the minimum number of samples in each condition. Using EdgeR, log2-fold changes in the gene expression between all condition pairs were calculated and further used for running gene set enrichment analysis.

#### Gene set enrichment analysis

The distribution of genes from KEGG pathways (http://www.kegg.jp) ^24^ in ranked gene lists was examined using gene set enrichment analysis (GSEA) ^25^. Ranked gene lists were formed based on log2-fold changes in the gene expression of Huh-7 cells cultured on a given basement membrane matrix in comparison to traditional polystyrene coated cell culture plates. The enrichment was considered to be significant if the corresponding false discovery rate (FDR) was smaller than 0.05. For visualizing GSEA results we considered both FDR and normalized enrichment score (NES) indicating if the enrichment was based on up-regulated (NES>0) or down-regulated (NES<0) genes (see Figure 3A). For the hierarchical clustering of KEGG pathways we used Euclidean distance for significance measures 1-FDR for pathways with NES>0 and FDR-1 for pathways with NES<0.

#### Estimating transcription factor activities

We used ISMARA ^26^ to estimate the activity of all transcription factors (TFs) with known binding specificity in both traditional and ECM-based Huh-7 cell cultures. In brief, ISMARA models the expression levels of all mRNAs in a sample in terms of the activity of all TFs as well as the response of respective gene promoters. The response of a promoter to a TF is assumed to be the number of predicted binding sites the promoter has for the respective TF. The explanatory power of any given regulatory motif in the data set is then represented by its z-score.

## Results and Discussion

To validate the morphological phenotypes reported earlier for the growth of Huh-7 cells on Matrigel basement membrane matrices ^14^ and compare them to those observed on Collagen matrices of different concentrations, we imaged the growing cells at Day 1 and Day 7 of seeding as mentioned in the methods section (Figure 1). Huh-7 cells grown on basement membrane matrices demonstrated a strikingly different morphology in comparison to cells grown on traditional polystyrene coated cell culture plates (Figure 1). To identify core transcriptional pathways that underlie the observed differences, RNA-seq samples from Huh-7 cells grown for 7 days on polystyrene coated cell culture plates (Control) and Matrigel (1.5 mg/ml and 3.0 mg/ml) and Collagen (0.5 mg/ml and 1.0 mg/ml) basement membrane matrices were generated from three biological replicates per condition. The resulting RNA-Seq data set was subjected to principal component analysis (PCA) (Figure 2A,B). Strikingly, the first principal component (PC1), which explained about 75% of the variance present in the data set, distinguished cells cultured on the Collagen matrix from cells cultured in other conditions. In contrast, PC2 reflected the increase in concentration of matrix components (Fig. 2A). To identify the molecular changes that contributed most to the variance explained by PC1 and PC2, we aligned the vectors given by the gene expression levels of individual genes across all samples with PC1 and PC2 (Figure 3) and subjected obtained genes to the gene ontology (GO) analysis with DAVID ^21^. Genes aligned with PC1 corresponded to GO terms such as “extracellular region/space”, “calcium ion binding”, “regulation of cell growth”, “regulation of cellular amino acid metabolic process” and “proteasome complex” (Figure 2C). Genes aligned with PC2 corresponded to the GO terms “extracellular region/space”, “inflammatory response” and “plasma membrane” (Figure 2D). Note that although some of the GO terms that were enriched by genes aligned with PC1 and PC2 were similar, the enrichment was due to distinct sets of genes (see Figures S2 and S3).

**Figure 1.**
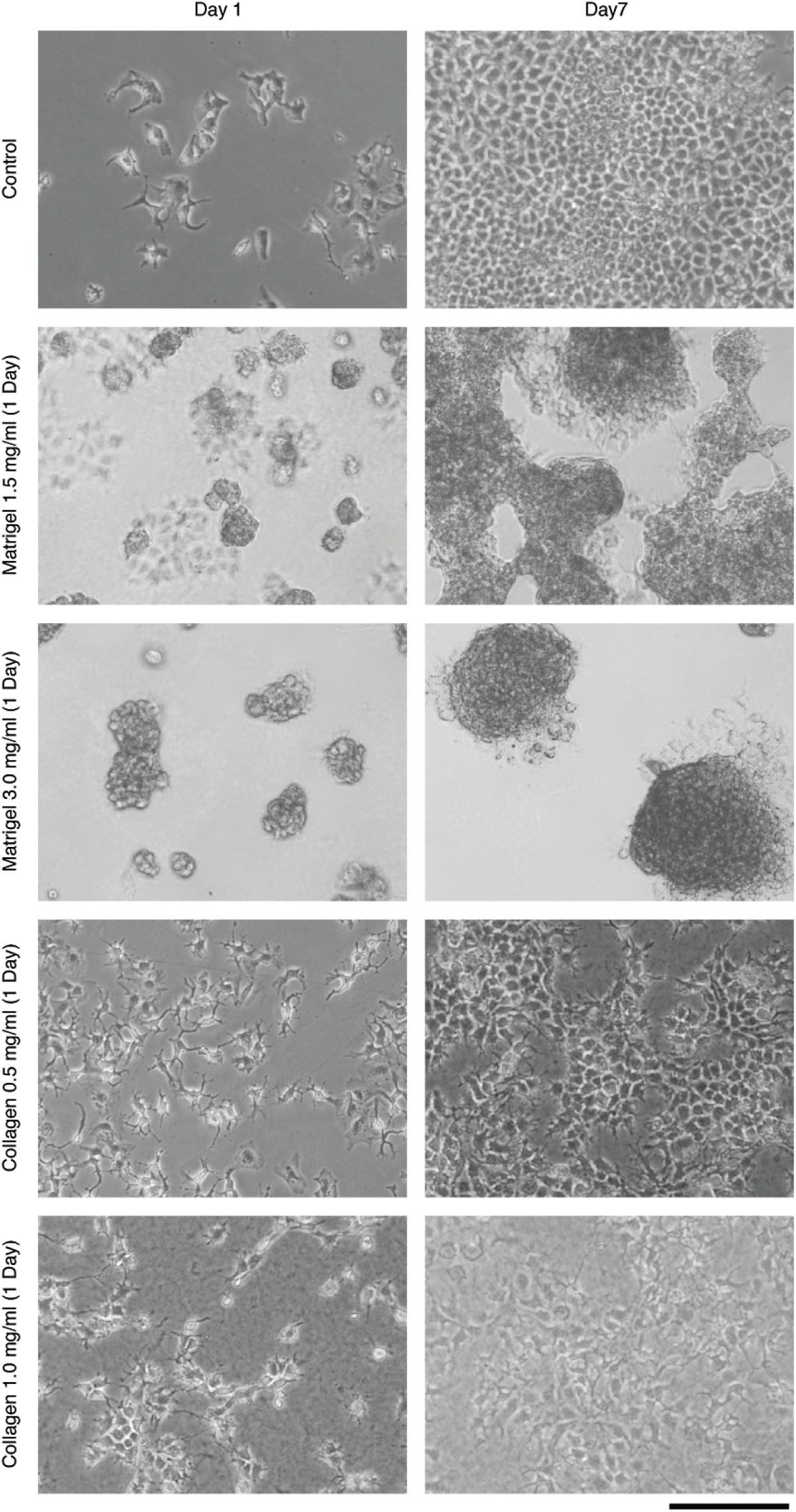
Phase contrast images of Huh-7 cells on the indicated basement membrane matrices or control culture medium at day 1 and day 7 post seeding. Scale bar marked below represents 500 μm.

**Figure 2.**
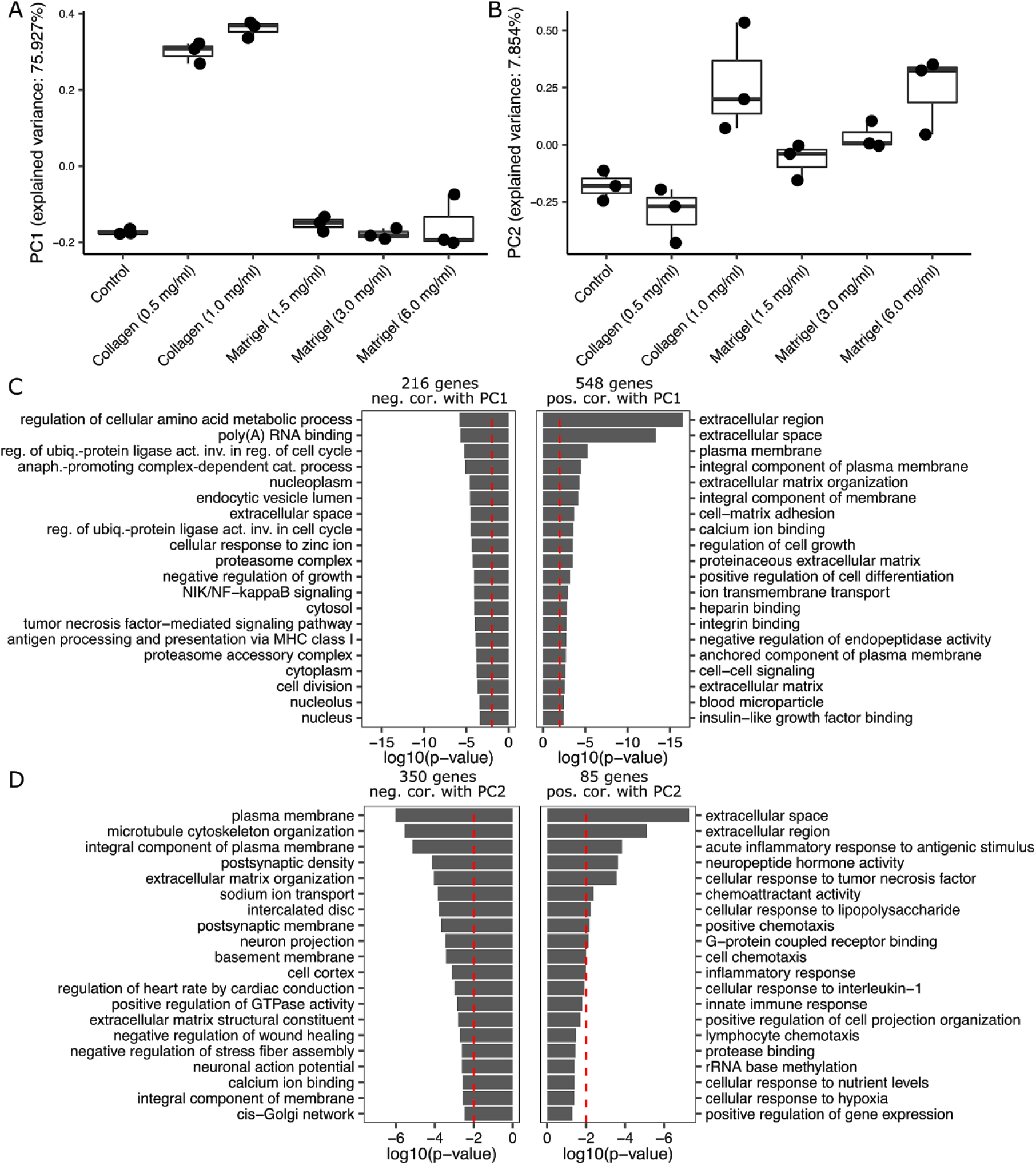
Principal component analysis (PCA) of the RNA-Seq data set prepared for samples of Huh-7 cells grown for 7 days on polystyrene coated cell culture plates (Control), or on Matrigel (1.5 mg/ml and 3.0 mg/ml), or on Collagen (0.5 mg/ml and 1.0 mg/ml) basement membrane matrices. A) Coordinates of PC1. B) Coordinates of PC2. Each dot corresponds to one sample. Samples are grouped by conditions. The numbers associated with the principal components indicate the fraction of the variance in transcript expression in that is explained by the corresponding principal component. C) Gene ontology (GO) analysis of genes aligned with PC1. D) GO analysis of genes aligned with PC2. GO analysis was performed with DAVID ^21^. Numbers on the top of bar plots indicate the number of genes used for the GO analysis. Top 20 enriched GO terms were visualized. As significance threshold for the enrichment we considered p-value<0.01 (dashed red lines).

**Figure 3.**
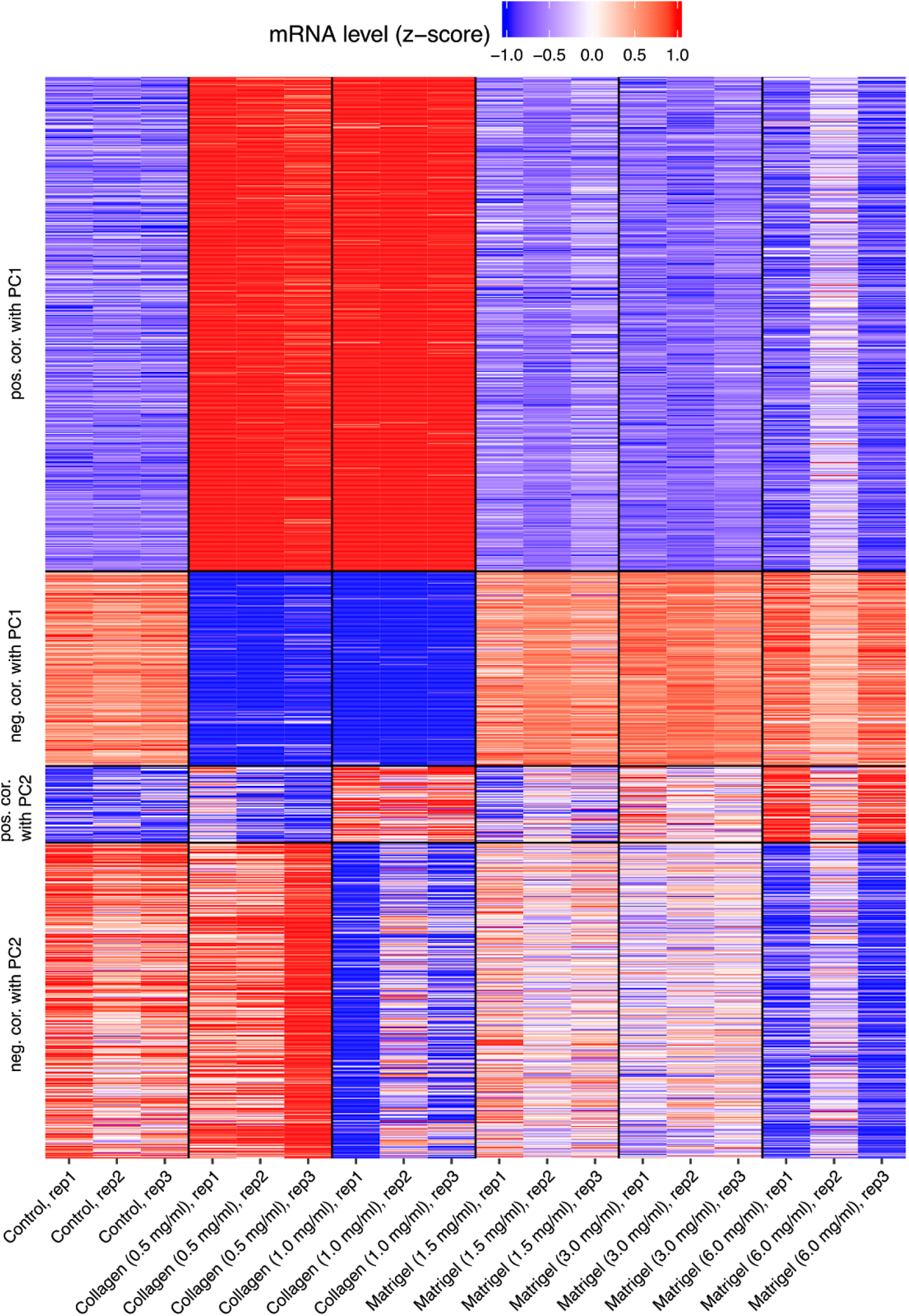
Heatmap of genes aligned with PC1 and PC2 and contributing most to the variance in the RNA-Seq data that is explained by these PCs.

We further carried out a gene set enrichment analysis (GSEA) ^25^ to determine which KEGG pathways are involved in the response of cells cultured on basement membrane matrices relative to traditional culture conditions (Figure 4A). This analysis revealed pathways that responded similarly to the ECM conditions, e.g. “Insulin secretion”, “Biosynthesis of amino acids”, “p53 signaling pathway” and “Ubiquitin mediated proteolysis”, as well as pathways that responded in opposite manner in cells grown on the collagen matrix in comparison to cells grown on matrigel, e.g. “ECM-receptor interaction”, “Focal adhesion” and “Cortisol synthesis and secretion”. There were also multiple pathways regulated only in one of the two growth conditions. For example, pathways related to DNA and RNA processing and homeostasis such as “Mismatch repair”, “mRNA surveillance pathway” and “RNA transport” were specifically enriched in cells grown on the collagen basement membrane matrix. In contrast, “FoxO signaling pathway” and “ErbB signaling pathway” were specifically enriched in cells grown on the matrigel matrix.

**Figure 4.**
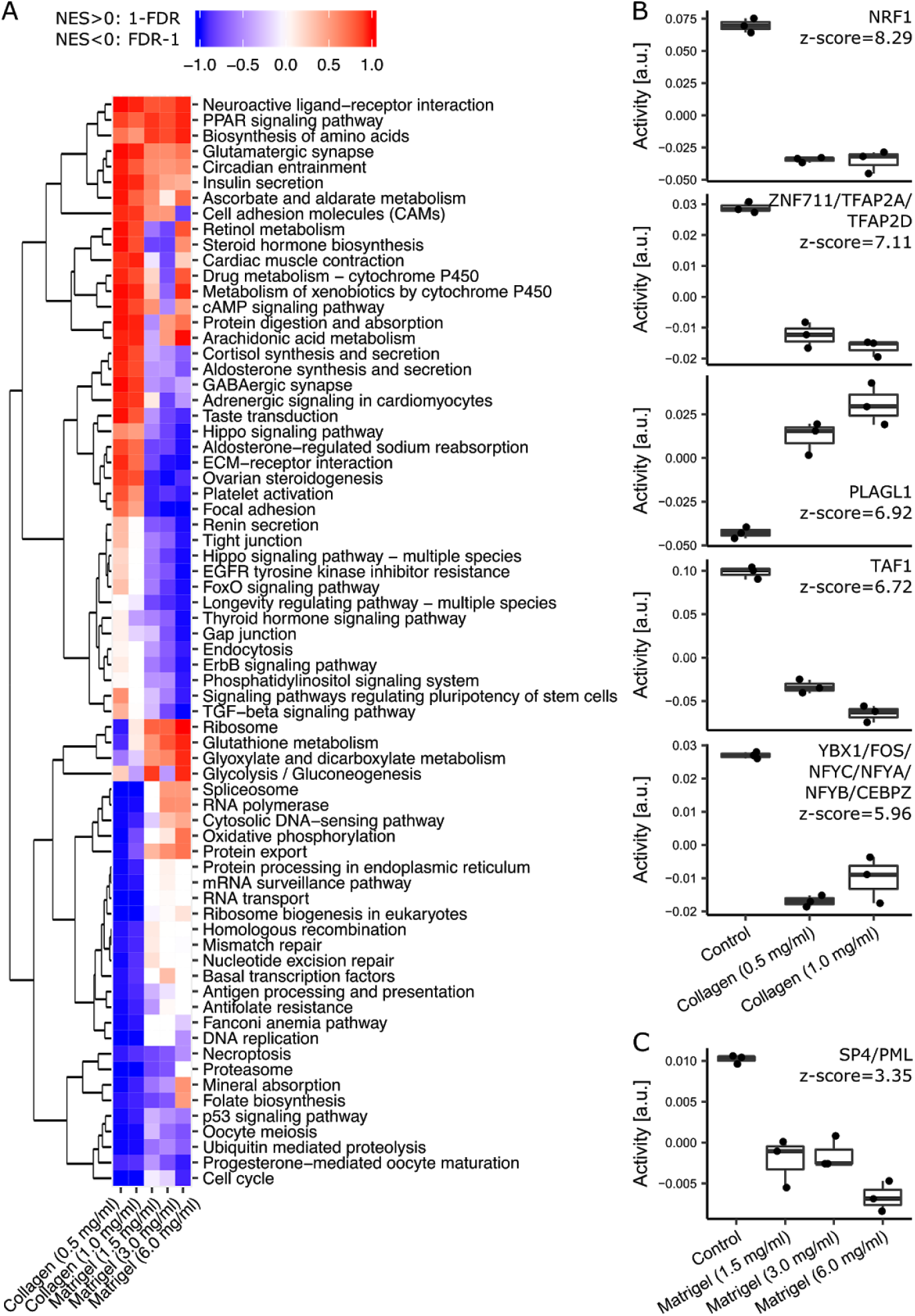
Pathway and motif activities in samples of Huh-7 cells grown for 7 days on polystyrene coated cell culture plates (Control), Matrigel (1.5 mg/ml and 3.0 mg/ml) or Collagen (0.5 mg/ml and 1.0 mg/ml) basement membrane matrices. A) Heatmap summarizing the enrichment of KEGG pathways among genes ranked by expression changes in cells grown on a basement membrane matrix in comparison to traditionally cultured cells. A pathway was included in the heatmap if it was significantly enriched for at least one condition. FDR<0.05 was chosen as significance threshold. B) Activities of the 5 most active motifs in collagen culture condition. C) Activity of the top motif in the matrigel culture condition. Each dot corresponds to one sample. Samples are grouped by conditions. Z-scores indicate the contribution of regulatory motifs associated with TF(s) to explaining gene expression changes.

Since numerous genes and pathways were regulated in Huh-7 cells upon changing the growth environment, we wondered which transcription factors (TFs) may be responsible for their regulation. To address this question, we estimated activities of TF binding sites across conditions using ISMARA ^26^. Figure 4B represents the inferred activities of the 5 most significant binding motifs and associated TFs from the comparison of control cells with cells grown on the collagen basement membrane matrix. Regulatory motifs whose activity decreased in cells grown on the collagen basement membrane matrix are associated with TFs NRF1 (targets annotated as “RNA polymerase”, “Chromosome organisation”), ZNF711/TFAP2A/TFAP2D (“Regulation of transferase activity”), TAF1 (“Spliceosome”, “RNA processing”) and YBX1/FOS/NFYC/NFYA/NFYB/CEBPZ (“Chromatin silencing”, “Protein folding” and “Cell cycle”). The regulatory motif whose activity increased in cells grown on the collagen basement membrane matrix is associated with the TF PLAGL1 (targets annotated as “Actin cytoskeleton”). Gene expression changes were milder in the Matrigel culture conditions and consequently, few TF showed consistently changed activity in this condition. The activity of the binding motif associated with the TF SP4/PML (targets annotated as “ECM”) that contributed most to the gene expression changes in Matrigel compared to control culture conditions and showed consistency between replicates is depicted in Figure 4C.

Our study reveals large differences in gene expression attributable to the specific culture conditions of Huh-7 and it highlights transcription factors that seem to underlie these differences. We believe that these data will aid the design of further mechanistic studies geared towards the development of an *in vivo* model for the study of liver function and disease.

**Online Table 1 :** Experimental study with replicates.

| Source | Protocol 1 | Protocol 2 | Samples | Protocol 3 | Data |
| --- | --- | --- | --- | --- | --- |
| Huh7 cells | Cell culture | RNA extraction | Control (Sample 1) | RNA Sequencing | GSM4222657 |
| Huh7 cells | Cell culture | RNA extraction | Control (Sample 2) | RNA Sequencing | GSM4222658 |
| Huh7 cells | Cell culture | RNA extraction | Control (Sample 3) | RNA Sequencing | GSM4222659 |
| Huh7 cells | Cell culture | RNA extraction | 3D Collagen (0.5 mg/ml) (Sample 1) | RNA Sequencing | GSM4222660 |
| Huh7 cells | Cell culture | RNA extraction | 3D Collagen (0.5 mg/ml) (Sample 2) | RNA Sequencing | GSM4222661 |
| Huh7 cells | Cell culture | RNA extraction | 3D Collagen (0.5 mg/ml) (Sample 3) | RNA Sequencing | GSM4222662 |
| Huh7 cells | Cell culture | RNA extraction | 3D Collagen (1.0 mg/ml) (Sample 1) | RNA Sequencing | GSM4222663 |
| Huh7 cells | Cell culture | RNA extraction | 3D Collagen (1.0 mg/ml) (Sample 2) | RNA Sequencing | GSM4222664 |
| Huh7 cells | Cell culture | RNA extraction | 3D Collagen (1.0 mg/ml) (Sample 3) | RNA Sequencing | GSM4222665 |
| Huh7 cells | Cell culture | RNA extraction | 3D Matrigel (1.5 mg/ml) (Sample 1) | RNA Sequencing | GSM4222666 |
| Huh7 cells | Cell culture | RNA extraction | 3D Matrigel (1.5 mg/ml) (Sample 2) | RNA Sequencing | GSM4222667 |
| Huh7 cells | Cell culture | RNA extraction | 3D Matrigel (1.5 mg/ml) (Sample 3) | RNA Sequencing | GSM4222668 |
| Huh7 cells | Cell culture | RNA extraction | 3D Matrigel (3.0 mg/ml) (Sample 1) | RNA Sequencing | GSM4222669 |
| Huh7 cells | Cell culture | RNA extraction | 3D Matrigel (3.0 mg/ml) (Sample 2) | RNA Sequencing | GSM4222670 |
| Huh7 cells | Cell culture | RNA extraction | 3D Matrigel (3.0 mg/ml) (Sample 3) | RNA Sequencing | GSM4222671 |
| Huh7 cells | Cell culture | RNA extraction | 3D Matrigel (6.0 mg/ml) (Sample 1) | RNA Sequencing | GSM4222672 |
| Huh7 cells | Cell culture | RNA extraction | 3D Matrigel (6.0 mg/ml) (Sample 2) | RNA Sequencing | GSM4222673 |
| Huh7 cells | Cell culture | RNA extraction | 3D Matrigel (6.0 mg/ml) (Sample 3) | RNA Sequencing | GSM4222674 |

---

**Figure S1.**
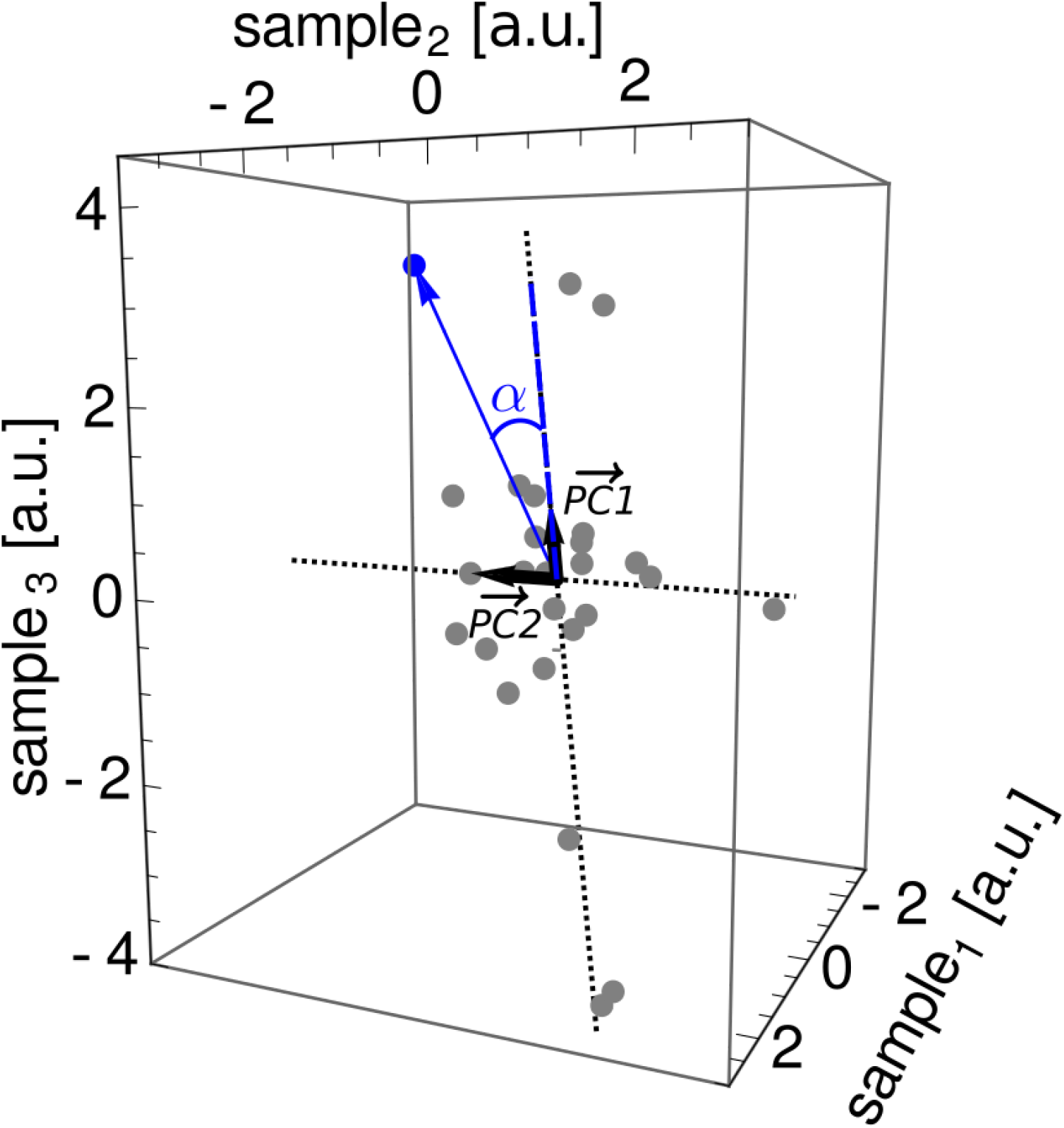
Visualization of the projection (blue dashed line) of a representative gene vector (blue arrow) on PC1 (black arrow) with the corresponding angle α between the gene vector and PC1. The correlation between the gene vector and PC1 corresponds to cos α.

**Figure S2.**
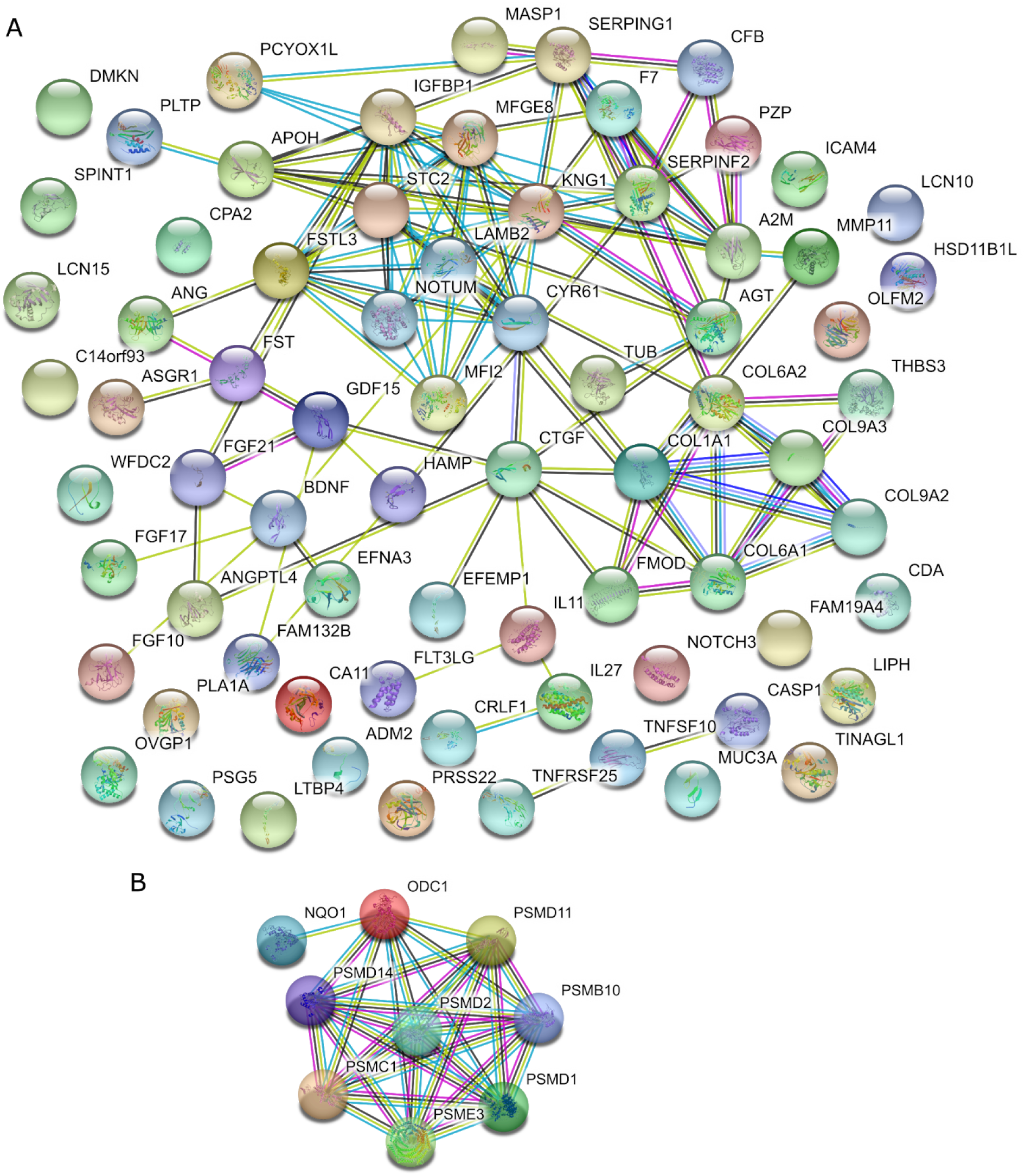
STRING networks of genes aligned with PC1 and associated with the top enriched GO categories. A) Genes associated with the GO term “Extracellular region”. B) Genes associated with the GO term “Regulation of cellular amino acid metabolic process”.

**Figure S3.**
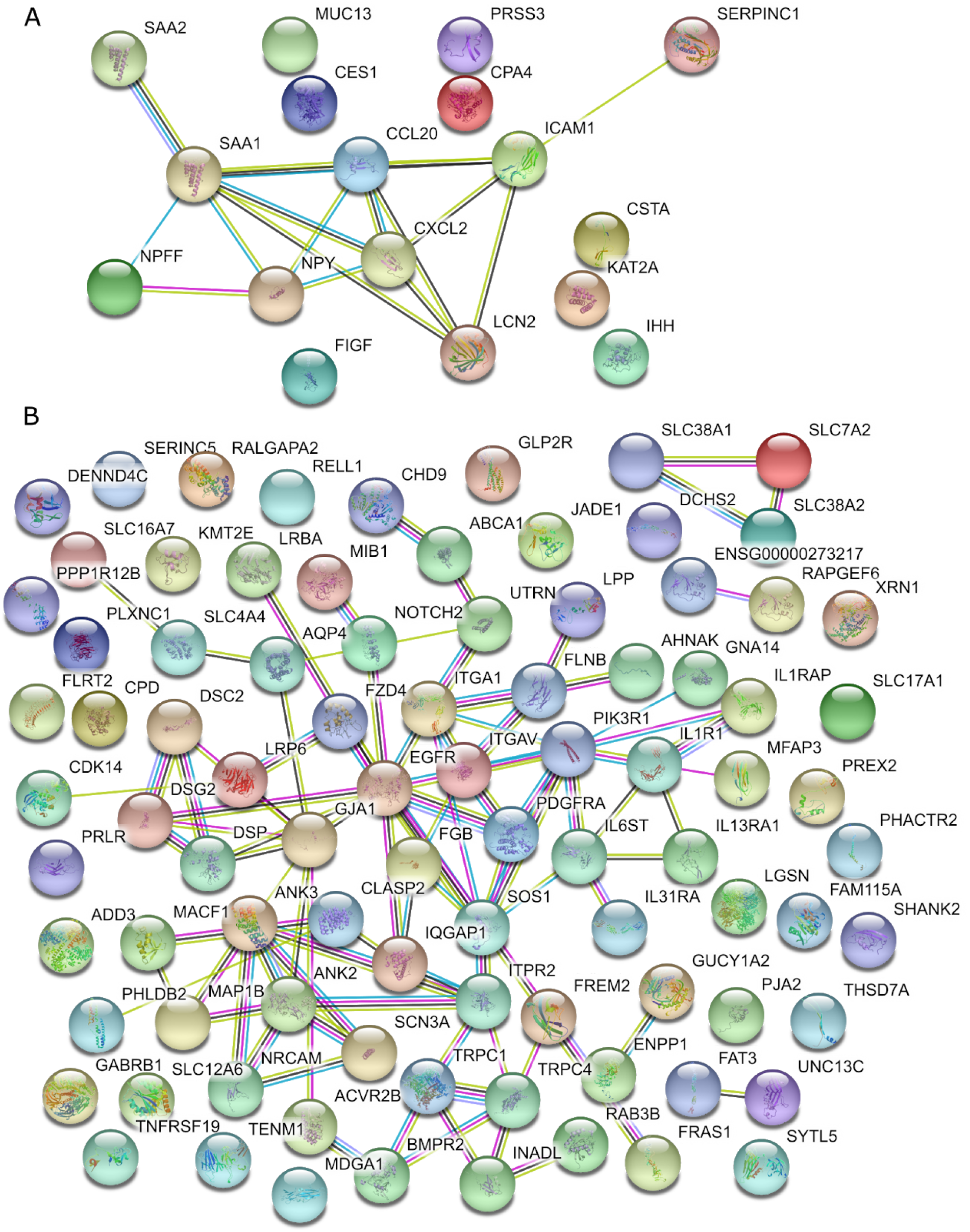
STRING networks of genes aligned with PC2 and associated with the top enriched GO categories. A) Genes associated with the GO term “Extracellular space”. B) Genes associated with the GO term “Plasma membrane”.

## Data Records

The RNA-seq data are available in the GEO database ^27,28^ as “ The transcriptional landscapes of a hepatoma cell line grown on scaffolds of Extracellular Matrix proteins ”, accession GSE142206.

## Code Availability

The code for the analysis of RNA-Seq data presented in the manuscript is available on GitHub: https://github.com/zavolanlab/Liver3DCulture.

## Acknowledgements

We would like to thank Philippe Demougin from the Genomics Facility Basel for genomic library preparation, and the sciCORE team for their maintenance of the HPC facility at the University of Basel. We also acknowledge intramural research funding for support.

## Author contributions

Conceptualization, design and execution of experiments - Souvik Ghosh

RNA seq data analysis - Anastasiya Börsch

Manuscript preparation - Souvik Ghosh and Anastasiya Börsch

Supervision of study and manuscript writing - Mihaela Zavolan

## Competing interests

The authors declare that there is no conflict of interest.

## Notes

### Competing Interest Statement

The authors have declared no competing interest.

